# Autonomic neurons are intrinsically hyperexcitable in a Friedreich ataxia iPSC model

**DOI:** 10.64898/2026.10.06.757136

**Authors:** Jarmon G. Lees, Haoxiang Zhang, Li Li, Mirella Dottori, Alice Pébay, Louise Corben, Martin Delatycki, Stephen Wilcox, Jeffrey M. Pullin, Jarny Choi, Davis McCarthy, Jill S. Napierala, Marek Napierala, Shiang Y. Lim

## Abstract

Friedreich ataxia (FRDA) is an inherited multisystem disorder caused by a GAA repeat expansion in the *FXN* gene and consequent frataxin deficiency, causing progressive neurodegeneration and heart disease. Clinical studies have documented autonomic dysfunction in FRDA, including elevated resting heart rate, raised plasma catecholamines and a high burden of urinary, bowel and sexual symptoms. Consistent with this, autopsy studies show degeneration of the cardiac conduction system and its associated ganglia in FRDA. Whether these features are intrinsic to FRDA autonomic neurons has been untestable without a human cellular model. Here, a human model of FRDA autonomic neurons was established by differentiating induced pluripotent stem cells (iPSCs) from individuals with FRDA and from their CRISPR/Cas9-corrected isogenic controls. FRDA autonomic neurons expressed canonical autonomic lineage markers and retained reduced *FXN* mRNA and frataxin protein levels. Electrophysiological analysis revealed pronounced hyperexcitability, with an approximately five-fold increase in spike number, shortened interspike intervals, and longer bursts. FRDA autonomic neurons also exhibited increased cell death, elevated mitochondrial reactive oxygen species, and a shift from networked to tubular mitochondrial morphology. Bulk RNA sequencing of all three isogenic pairs revealed loss of extracellular matrix programs and activation of stress response pathways. The top differentially expressed gene was *MEG3*, a maternally imprinted long noncoding RNA, which was consistently upregulated across all three isogenic pairs. *MEG3* sits within the imprinted 14q32 DLK1–DIO3 locus, whose maternally expressed noncoding transcripts were co-ordinately de-repressed in FRDA autonomic neurons. Competitive gene-set testing of 43 imprinted genes across 20 domains showed this to be specific to 14q32: no other imprinted domain was co-ordinately altered. Together, these findings establish hyperexcitability of autonomic neurons as a cell autonomous consequence of frataxin deficiency, accompanied by mitochondrial dysfunction. This provides a cellular basis for the increased sympathetic tone observed in clinical studies.

## Introduction

Friedreich ataxia (FRDA) is one of the most common inherited ataxias, characterised by progressive neurodegeneration and multisystem involvement, primarily affecting the nervous system and the heart [23, 52]. Symptom onset typically occurs between ages 8 and 15, and neurological manifestations include progressive limb and gait ataxia, dysarthria and sensory loss, while non-neurological features include scoliosis, diabetes and cardiomyopathy [17, 23, 34]. Cardiac complications, particularly arrhythmias and congestive heart failure, are the leading causes of mortality, accounting for approximately 60% of deaths, with around one in eight deaths attributable to arrhythmia [17, 35, 38]. Omaveloxolone is the only approved therapy, and none targets the cardiac phenotype specifically [28, 42, 47].

FRDA is caused by mutations in *FXN* on chromosome 9, which encodes the mitochondrial protein frataxin. Approximately 96% of affected individuals are homozygous for a GAA trinucleotide repeat expansion in intron 1, which induces repressive heterochromatin and suppresses *FXN* transcription [3, 34, 52]. Frataxin is required for iron–sulfur cluster biosynthesis, and its deficiency disrupts mitochondrial iron homeostasis, leading to mitochondrial dysfunction, oxidative stress, and progressive damage to nervous and cardiovascular tissues [12, 29].

The cardiac autonomic nervous system has been implicated in the pathogenesis of FRDA cardiomyopathy [29]. Post-mortem studies describe focal degeneration and fibrosis of the sinus node, together with loss of myelinated and unmyelinated nerves and ganglia within the conduction system [19, 20]. Clinical studies also point to autonomic involvement, with elevated plasma catecholamines [39, 46] and reports of dysautonomia with labile hypertension in individuals with FRDA [37]. Pelvic autonomic symptoms are also common, with 80% of FRDA individuals in one cohort reporting lower urinary tract symptoms, 64% bowel symptoms and 83% sexual symptoms [6, 25], and the European registry recorded urinary dysfunction in 42.8% of 650 individuals with FRDA [51].

Notably, formal autonomic function testing shows little evidence of autonomic failure. Baroreflex sensitivity, Valsalva responses and sudomotor function are largely preserved, and the consistent abnormality across studies is an elevated heart rate at rest and during orthostatic challenge [16, 18]. This pattern has been interpreted as increased sympathetic tone rather than autonomic denervation [29]. Clinical measurements alone, however, cannot distinguish a defect intrinsic to autonomic neurons from a secondary response to cardiac disease.

Induced pluripotent stem cells (iPSCs) from individuals with FRDA provide a genetically accurate platform to resolve these questions. FRDA iPSCs have been differentiated into cardiomyocytes [7, 30], sensory and proprioceptive neurons [8, 9, 14, 58], cortical neurons [5, 11], and more recently into cerebellar organoids [53] and models containing microglia [49]. Protocols now also exist to derive autonomic neurons with postganglionic sympathetic character from human pluripotent stem cells [10, 55], and these neurons have revealed disease-relevant functional phenotypes in long QT syndrome type 1 and familial dysautonomia [54, 60]. To our knowledge, however, autonomic neurons have not been generated from FRDA iPSCs, and the use of CRISPR/Cas9-corrected isogenic controls provides a genetically matched comparison that helps to limit the confounding effect of genetic background and helps to identify the effects of frataxin deficiency.

In this study, autonomic neurons were differentiated from iPSCs derived from individuals with FRDA and their CRISPR/Cas9-corrected isogenic controls. Their marker expression, spontaneous electrical activity, mitochondrial function and viability were characterised, and the transcriptome of three isogenic pairs was profiled. The aims were to determine whether frataxin deficiency produces a cell-autonomous autonomic neuronal phenotype and to identify candidate molecular targets for translational follow-up.

## Materials and methods

### Human iPSC lines

Fibroblasts from three genetically confirmed adults with FRDA were reprogrammed into iPSC lines (FA1-3), and corresponding CRISPR/Cas9-corrected isogenic control lines (FA1ic-3ic) were generated as previously described [30, 33]. These iPSC lines were originally derived and characterised by Marek Napierala’s group (University of Texas Southwestern Medical Center). All iPSC lines were authenticated by Short Tandem Repeat analysis across 18 loci, with all edited lines matching 100% to their parental lines [40]. Our laboratory received the same authenticated stocks, and all cultures were routinely tested and confirmed mycoplasma free. Both FRDA and isogenic control iPSC lines were maintained on vitronectin (Thermo Fisher Scientific, MA, USA) coated plates and cultured in StemFlex medium (Thermo Fisher Scientific).

### Autonomic neuron differentiation

Autonomic neurons were derived from human iPSCs as previously described [24, 27, 30]. Briefly, FRDA and isogenic control iPSCs were seeded into ultra-low attachment round-bottom 96-well plates at a density of 2.0×10^4^ cells/well in E6 medium (Thermo Fisher Scientific) supplemented with 2 µM CHIR99021, 10 µM SB431542 (STEMCELL Technologies, Vancouver, Canada), and 10 µM Y-27632 (Tocris Bioscience, Bristol, UK) (designated as day 0). On day 3, the medium was replaced with E6 medium supplemented with 20 ng/mL FGF-2 (PeproTech, NJ, USA), 50 ng/mL

BMP4 (STEMCELL Technologies), 100 nM retinoic acid (Sigma-Aldrich, MA, USA) and 50 U/mL penicillin/streptomycin (Thermo Fisher Scientific). The medium was refreshed on day 7. On day 10, neurospheres were collected and dissociated with TrypLE™ Select Enzyme (Thermo Fisher Scientific), then re-seeded into 6-well ultra-low attachment plate (Corning, NY, USA) in Neurobasal Plus basal medium (Thermo Fisher Scientific) supplemented with N-2 Plus supplement (Thermo Fisher Scientific), B-27 Plus supplement (Thermo Fisher Scientific), GlutaMAX (Thermo Fisher Scientific), 20 ng/mL bFGF (PeproTech), 20 ng/mL EGF (PeproTech), 50 ng/mL BMP4, and 2 µg/mL heparin (Sigma-Aldrich). The medium was refreshed on day 14. Beginning on day 18, cultures were maintained in Neurobasal Plus medium supplemented with N-2 Plus supplement, B-27 Plus supplement, 10 ng/mL BDNF (PeproTech), 10 ng/mL GDNF (PeproTech), and 10 ng/mL NGF (PeproTech), with medium changes every 3-4 days to support neuronal maturation. On day 32, neurospheres were dissociated and either cryopreserved, or plated at 1.0×10^5^ cells/cm^2^ on Matrigel-coated tissue culture plates or multielectrode arrays in glucose-free DMEM medium supplemented with 5.55 mM glucose (Thermo Fisher Scientific), 10% foetal calf serum (Bovogen Biological, VIC, Australia), 10 ng/mL BDNF (PeproTech), 10 ng/mL GDNF (PeproTech), 10 ng/mL NGF (PeproTech), and 10 µM Y27632 (Tocris Bioscience). The medium was refreshed every 2-3 days without Y27632. On day 37, neurons were assayed for cellular function, mitochondrial function or RNA expression, or maintained on multielectrode arrays for 4 weeks, until day 60, for electrophysiological recording.

### Reverse transcription-quantitative polymerase chain reaction (RT-qPCR)

Total RNA was extracted from autonomic neurons using Tri-Reagent (Thermo Fisher Scientific), followed by RNA precipitation with chloroform and isopropanol (Sigma-Aldrich). cDNA was synthesized using a high-capacity cDNA reverse transcription kit (Applied Biosystems, CA, USA) [30]. Quantitative PCR was performed in duplicate using TaqMan Universal Master Mix, the QuantStudio 6 Flex Real-Time PCR system, and TaqMan Gene Expression Assays for *FXN* (Hs00175940_m1), *MEG3* (Hs00292028_m1), *MEG8* (Hs00419701_m1), *MEG9* (Hs01593046_s1), *DLK1* (Hs00171584_m1), *RTL1* (Hs06637009_s1), *DIO3* (Hs00956431_s1), and *GAPDH* (Hs99999905_m1) (Applied Biosystems, CT, USA). Relative mRNA expression was quantified using the comparative CT method (2^-ΔΔCt^) method, normalized to *GAPDH*.

### Frataxin protein expression

Relative frataxin protein expression was determined using the Frataxin Protein Quantity Dipstick Assay Kit (ab109881, Abcam) according to the manufacturer’s protocols [30]. Briefly, 4 µg of total protein from autonomic neurons, quantified using the Pierce™ BCA Assay (Thermo Fisher Scientific), was applied to microplate wells pre-coated with dried gold-conjugated frataxin antibody and incubated for 5 minutes at room temperature. The samples were then wicked onto dipsticks, allowed to dry at room temperature, and imaged using a flatbed scanner. Band intensities were quantified using ImageJ software.

### Immunofluorescence

Autonomic neurons were fixed in 10% neutral buffered formalin (Trajan Scientific, VIC, Australia) for 20 minutes at room temperature and permeabilized with 0.2% Triton X-100 for 10 minutes. After blocking with serum-free blocking solution (Dako, VIC, Australia) for 10 minutes, cells were stained with PHOX2B (2 µg/mL, mouse monoclonal, sc376997, Santa Cruz), tyrosine hydroxylase (5 µg/mL, rabbit monoclonal, ab137869, Abcam), MASH1 (5 µg/mL, rabbit monoclonal, ab211327, Abcam), PRPH (2 µg/mL, mouse monoclonal, sc377093, Sanat Cruz), or Hsp60 (2 µg/mL, rabbit polyclonal, ab46798, Abcam) primary antibodies overnight at 4°C, followed by secondary antibodies Alexa Fluor 488-conjugated goat-anti-rabbit or goat-anti-mouse (10 μg/mL, A11001/A11008, Invitrogen), and Alexa Fluor 594-conjugated goat-anti-mouse or goat-anti-rabbit (10 μg/mL, A11005/A11012, Invitrogen) for 1 hour at room temperature. Cells were then counterstained with 1 μg/mL of DAPI (Invitrogen) and mounted in fluorescence mounting medium (Dako),

#### Mitochondrial morphology

Mitochondrial morphology was assessed by Hsp60 immunostaining of autonomic neurons as previously described [13, 30]. Fixed cells were stained with PHOX2B and Hsp60, and images were acquired using an Olympus BX61 fluorescence microscope at 600x magnification. For each replicate, images were captured from 10 randomly selected fields of view, and a minimum of 60 PHOX2B-positive autonomic neurons were analysed. Mitochondrial morphology was classified as either predominantly networked or tubular, based on whether more than 50% of the mitochondria in a given cell exhibited that specific morphology. For each condition, the percentages of cells classified as having networked or tubular mitochondria were calculated, summing to 100%.

### Electrophysiological assessment

Autonomic neurons were seeded onto a 60-electrode two-dimensional microelectrode array (60-MEA200/12/80iR-Ti-gr, Multichannel Systems, Germany) and allowed to mature for 28 days in neural maintenance medium. Recordings were then performed using an MEA1060-2BC system (Multichannel Systems) with the array maintained at 37 °C. Recordings were acquired at a sampling rate of 25 kHz and band-pass filtered between 300 and 6000 Hz using a second-order Butterworth filter. Five minutes of spontaneous activity were recorded per experiment, and values from all active channels of the 60-channel array were averaged to give a single value per recording. Spikes were detected on the filtered signal using an amplitude threshold of 4σ_n, where σ_n = median(|x| / 0.6745) and x is the filtered signal [50]. This median-based estimate of the background noise standard deviation is robust to the presence of spikes within the segment used for its calculation, and was derived from a 500 ms window of filtered signal. Bursts were detected in MC_Rack (Multichannel Systems) using the MaxInterval method [32]. A burst was initiated when an interspike interval of ≤ 100 ms was encountered and continued while intervals remained ≤ 100 ms, ending at the first interval exceeding this value. Bursts separated by less than 200 ms were merged, and bursts of less than 10 ms duration or containing fewer than 4 spikes were discarded.

### Cell death assay

Autonomic neuron viability was assessed using propidium iodide and Hoechst 33258 staining. Cells were incubated with 5 μg/mL of propidium iodide and 1 μg/mL of Hoechst 33258 (Sigma-Aldrich), prepared in HBSS++, for 45 minutes at 37°C in a humidified CO_2_ incubator, protected from light. Fluorescent images were captured using an inverted Olympus IX-71 microscope at 200x magnification. The number of dead cells (propidium iodide positive) was counted and expressed as a percentage of the total number of cells (Hoechst 33258 positive). At least 100 cells per condition were analysed in each independent experiment.

### Mitochondrial ROS measurement

Mitochondrial ROS levels in autonomic neurons were assessed using the MitoSOX™ Red dye (Thermo Fisher Scientific). Cells were incubated with 5 μM MitoSOX™ Red in HBSS++ for 20 minutes at 37°C in a humidified CO_2_ incubator, protected from light. Following incubation, cells were washed once with HBSS++ and imaged immediately in fresh HBSS++ using an inverted Olympus IX-71 microscope at 200x magnification. The corrected total cell fluorescence was quantified for 100 autonomic neurons from 3 random fields per group in each independent experiment.

### Mitochondrial membrane potential measurement

Mitochondrial membrane potential was assessed using tetramethylrhodamine methyl ester (TMRM, Thermo Fisher Scientific). Autonomic neurons were incubated with 10 nM TMRM in HBSS++ (non-quenching concentration) for 20 minutes at 37°C in a humidified CO_2_ incubator, protected from light. Imaging was performed immediately after incubation, without a wash step, using an inverted Olympus IX-71 microscope at 200x magnification. The corrected total cell fluorescence was quantified for 100 autonomic neurons across 3 random fields for each group in each independent experiment.

### RNA sequencing and analysis

Total RNA was extracted from autonomic neurons derived from all three FRDA and isogenic control iPSC lines across three independent experiments using the Cytiva Mini RNA Extraction Kit (Cytiva, MA, USA), following the manufacturer’s instructions. Residual genomic DNA was removed by treatment with TURBO DNase (Thermo Fisher Scientific). RNA integrity was assessed using the Agilent 2100 Bioanalyzer (Agilent Technologies).

Library preparation was performed using 100 ng of input RNA per sample with the TruSeq Stranded Total mRNA Library Prep Kit (poly A mRNA enrichment via oligo dT magnetic beads; Illumina, CA, USA) as per manufacturer’s instruction. The library was quantified using the Agilent Tapestation and the Qubit™ RNA assay kit for Qubit 2.0® Fluorometer (Life technologies). Unique dual indexes (Illumina TruSeq RNA UD Indexes) were used to barcode individual libraries. The indexed libraries were then prepared and diluted to 750pM for paired end (2x 101 base) sequencing on a NextSeq2000 instrument using the P3 200 cycle kit using v3 chemistry (Illumina, CA, USA) as per manufacturer’s instructions. The base calling and quality scoring were determined using Real-Time Analysis on board software v2.4.6, while the FASTQ file generation and de-multiplexing utilised bcl2fastq conversion software v2.15.0.4. Libraries yielded a median of 31.95 million read pairs per sample, with a median Q30 score of 87.9% and a median Salmon mapping rate of 87.6%.

RNA sequencing (RNA-seq) data were processed and analyzed using the ARMOR (Automated Reproducible Modular Workflow for Preprocessing and Differential Analysis of RNA-seq Data) pipeline [45]. Following read quality assessment and adapter trimming, adapters and low□quality bases were removed with Trim Galore! (□q 20). Transcript abundance was quantified using Salmon with quasi mapping to the Ensembl GRCh38 (release 93) transcriptome, with sequence□and GC□bias correction enabled. Gene-level counts were filtered with filterByExpr in edgeR, retaining 17,452 expressed genes (median 25,747 genes detected per sample), and normalised by the trimmed mean of M-values (TMM) method. Differential expression was assessed with edgeR quasi-likelihood F tests. Each FRDA line was compared with its matched isogenic control using a design containing disease status alone. For the combined analysis, patient of origin was included as a factor, so that the effect of frataxin deficiency was estimated within isogenic pairs and between-patient variation was excluded from the residual error. Genes were considered differentially expressed at a Benjamini–Hochberg FDR < 0.05.

For visualisation, principal component analysis was performed on the 2,000 most variable genes; the patient-of-origin effect was removed with removeBatchEffect (limma) using a design that retained disease status, and this adjusted matrix was used for display only and not for any statistical test. Variance attributable to patient and to disease was estimated by partitioning the per-gene sum of squares across all 17,452 expressed genes.

Up- and down-regulated genes were tested separately for overrepresentation by Fisher exact test against a background of the 17,071 unique gene symbols represented among the 17,452 expressed genes, using Gene Ontology Biological Process, Reactome [v97] and MSigDB Hallmark gene sets, with Benjamini–Hochberg correction. Near-duplicate terms were collapsed for display.

A candidate list of 56 imprinted genes across 21 domains was annotated with the normally expressed parental allele using the Geneimprint human catalogue (accessed September 2026), with four genes absent from that catalogue (*MEG9*, *SNHG14*, *ZNF331*, *TRAPPC9*) assigned from the published literature; 43 uniquely named genes across 20 domains were measurable among the 17,452 expressed genes. Nine gene sets were tested, in three families. Allele-specific sets comprised the maternally and paternally expressed genes, each divided into those within the 14q32 DLK1–DIO3 domain (n = 3 and 2) and those outside it (n = 11 and 24); this division distinguishes an effect confined to 14q32 from a general maternal- or paternal-allele phenomenon. Whole-imprintome sets comprised all 43 measurable imprinted genes (n = 43) and the same set excluding 14q32 (n = 38), testing whether imprinting was disturbed globally. Per-domain sets were formed for the three domains with at least five measurable genes: 14q32 DLK1–DIO3 (n = 5), 11p15.5 IGF2/H19 (n = 7) and 15q11–q13 PWS/AS (n = 7); the remaining 17 domains contributed three or fewer genes each and were not tested individually. Three genes with isoform-dependent imprinting (*GNAS*, *BLCAP*, *GRB10*) were excluded from the allele-specific sets. Sets were tested with camera, a competitive test accounting for inter-gene correlation, using the patient-blocked design (∼ patient + disease) on log-CPM values for all 17,452 genes, with Benjamini–Hochberg correction across the nine sets.

### Statistics

Sample size was predetermined by power calculation using a standard deviation of 10%, estimated from pilot experiments. A minimum of three independent experimental replicates provides >80% power to detect a difference of ≥22% using ANOVA (SD = 10%, α = 0.05, β = 0.2). Data are presented as mean ± standard error of the mean (SEM). Statistical significance was evaluated using two-sided unpaired Student’s t-test or Welch’s t-test when variances were unequal. For RNA sequencing, differential expression was assessed using edgeR with Benjamini–Hochberg false discovery rate (FDR) control (FDR < 0.05). Here, n denotes biological replicates (independent differentiations from distinct iPSC lines); technical replicates (multiple wells or imaging fields) were averaged within each biological replicate. Experiments were replicated in 3-6 independent runs unless otherwise specified, and exact n values are reported in the figure legends. Imaging fields were randomly selected for all analyses, and cells were randomly allocated to treatment conditions and to downstream assays. Assessors were blinded to experimental groups where possible. No data were excluded from analyses. A p-value < 0.05 was considered statistically significant.

## Results

### Differentiation of FRDA iPSCs into autonomic neurons

To establish a human cellular model of autonomic dysfunction in FRDA, autonomic neurons were differentiated from iPSCs of three individuals with FRDA (FA1–3) and their CRISPR/Cas9-corrected isogenic controls (FA1ic–3ic), in which the pathogenic GAA repeat expansion was excised. Both isogenic control and FRDA iPSC lines generated neurons that expressed the autonomic lineage transcription factors PHOX2B and MASH1 (ASCL1), together with tyrosine hydroxylase and peripherin, confirming a noradrenergic peripheral neuronal identity (Fig. 1a). *FXN* mRNA was reduced approximately 75% in FRDA autonomic neurons relative to isogenic control (Fig. 1b), with a corresponding ∼75% reduction in frataxin protein (Fig. 1c).

**Fig. 1.**
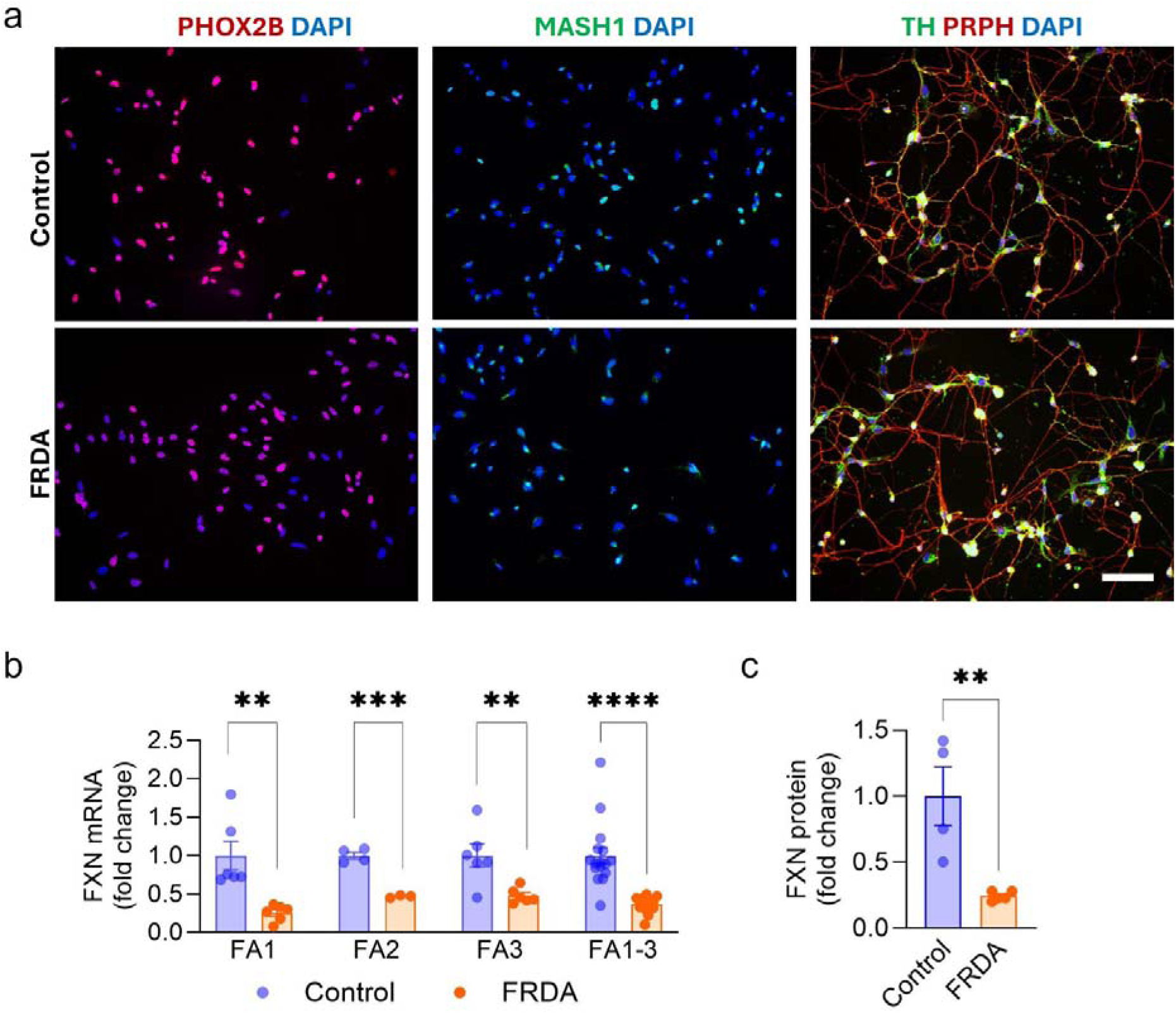
Generation and characterisation of autonomic neurons from Friedreich ataxia (FRDA) and isogenic control iPSCs. **a** Representative immunofluorescence images of autonomic neurons differentiated from FRDA (FA1) and isogenic control (FA1ic) iPSCs, stained for PHOX2B (red), MASH1 (green), tyrosine hydroxylase (TH, green) and peripherin (PRPH, red), with nuclei counterstained with DAPI (blue). Scale bar = 50 µm. **b** FXN mRNA in autonomic neurons from FA1, FA2, and FA3 iPSC lines, relative to matched isogenic controls. **c** FXN protein in the autonomic neurons from FA1 iPSC line and its isogenic control. Data are presented as mean ± SEM; n = 3-6 independent differentiations. **p < 0.01, ***p<0.001, ****p<0.0001 by unpaired Student’s t-test

### FRDA autonomic neurons are hyperexcitable

Spontaneous electrical activity was recorded by multielectrode array after 28 days of maturation (Fig. 2a). FRDA autonomic neurons showed an approximately five-fold increase in the maximum number of spikes (Fig. 2b) and a shorter mean interspike interval (Fig. 2c) relative to isogenic controls. This increase reflected a change in burst structure rather than burst frequency. The mean number of bursts was unchanged (Fig. 2d), while bursts in FRDA neurons contained more spikes (Fig. 2e) and lasted longer (Fig. 2f). FRDA autonomic neurons therefore fire longer, denser bursts at an unchanged burst rate (Fig. 2g), a pattern of sustained hyperexcitability.

**Fig. 2.**
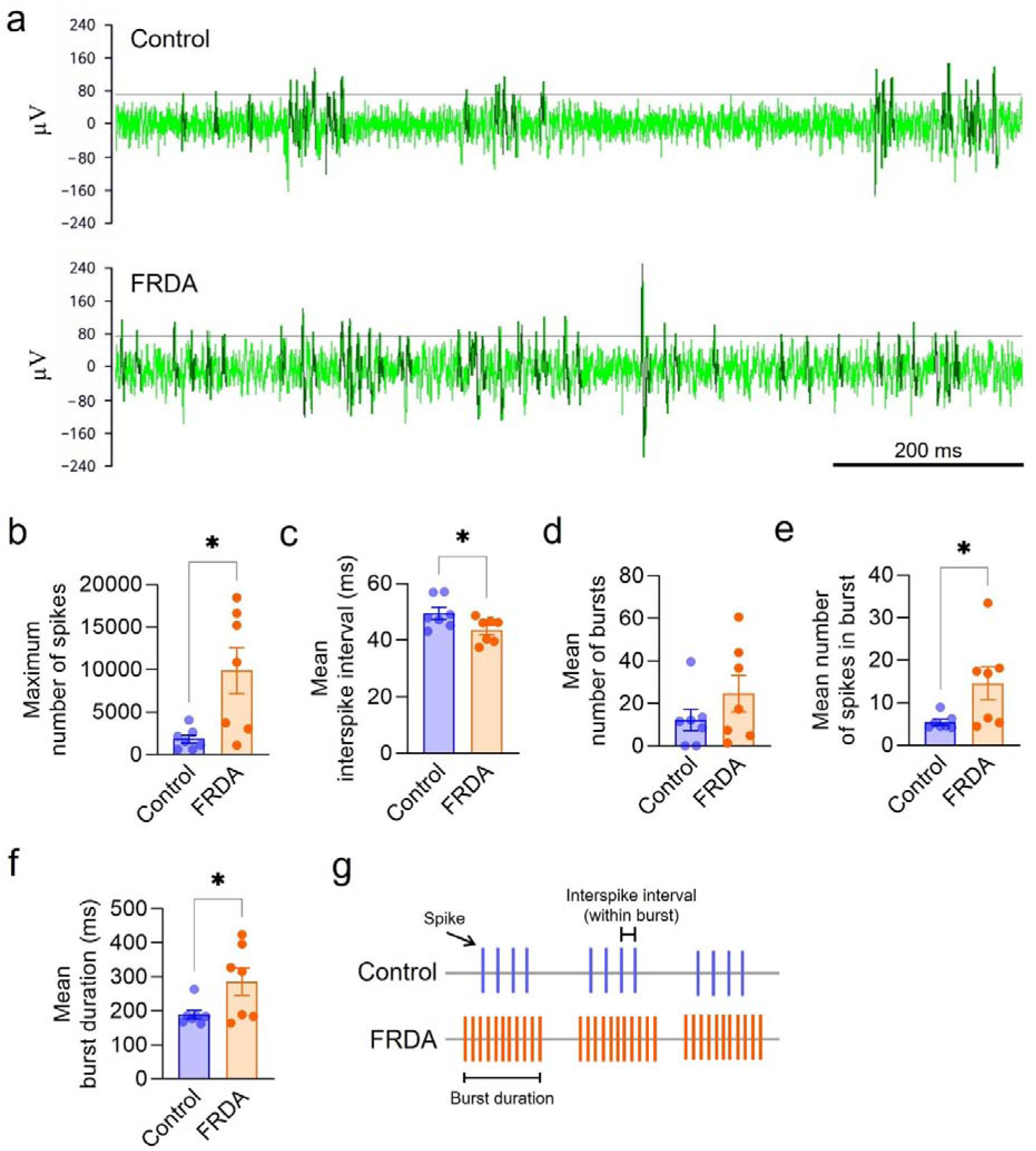
FRDA autonomic neurons are hyperexcitable. **a** Representative multielectrode array recordings from control and FRDA autonomic neurons after 28 days of maturation. **b-f** Quantification of **b** maximum number of spikes, **c** mean interspike interval, **d** mean number of bursts, **e** mean number of spikes per burst and **f** mean burst duration. **g** Schematic summarising the spiking pattern in control and FRDA autonomic neurons. Each data point represents one 5-minute recording, averaged across all active channels of the 60-electrode array. Data are presented as mean ± SEM; n = 4 independent differentiations. *p < 0.05 by unpaired Student’s t-test

### FRDA autonomic neurons show mitochondrial dysfunction and increased cell death

Frataxin is a mitochondrial protein essential for maintaining mitochondrial homeostasis and function, with critical implications for neuronal health. FRDA autonomic neurons exhibited approximately twice the level of cell death of isogenic controls (Fig. 3a) and elevated mitochondrial reactive oxygen species (Fig. 3b) relative to isogenic controls. Mitochondrial membrane potential was also reduced in FRDA autonomic neurons, though this did not reach statistical significance (Fig. 3c). Mitochondrial morphology was also altered, with FRDA neurons showing a lower proportion of networked mitochondria and a higher proportion of short tubular mitochondria (Fig. 3d), indicating a fragmented mitochondrial phenotype. To identify the transcriptional changes underlying these phenotypes, FRDA autonomic neurons were next profiled by RNA sequencing.

**Fig. 3.**
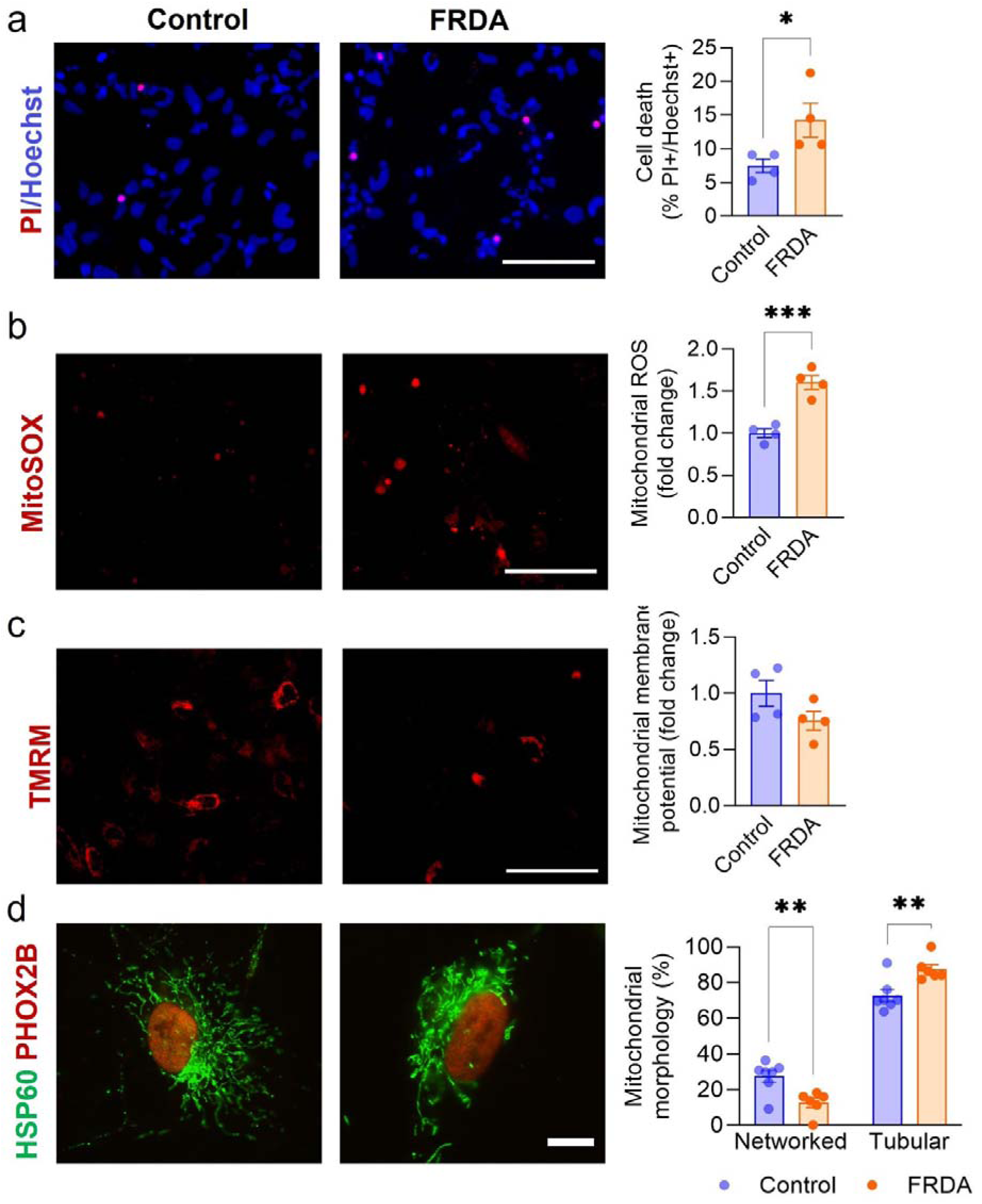
FRDA autonomic neurons show mitochondrial dysfunction and increased cell death. Each panel shows representative fluorescence images of control (left) and FRDA (right) autonomic neurons with the corresponding quantification at right. **a** Cell death assessed by propidium iodide (PI, red) uptake with Hoechst counterstain (blue), quantified as the percentage of PI-positive nuclei relative to total Hoechst-positive nuclei. **b** Mitochondrial reactive oxygen species visualised with MitoSOX Red, expressed as fold change. **c** Mitochondrial membrane potential visualised with TMRM, expressed as fold change. **d** Mitochondrial morphology classified as networked or tubular in neurons immunostained for HSP60 (green) and PHOX2B (red). Scale bar = 100 µm **a-c**, 10 µm **d**. Data are presented as mean ± SEM. n = 4-6 independent differentiations. *p < 0.05, **p < 0.01, ***p < 0.001 by unpaired Student’s t-test

### Transcriptomic profiling of FRDA autonomic neurons

To determine the transcriptional consequences of frataxin deficiency in autonomic neurons, bulk RNA sequencing was performed on autonomic neurons differentiated from three FRDA iPSC lines (FA1–FA3) and their CRISPR/Cas9-corrected isogenic controls (FA1ic–FA3ic), with three independent differentiations per cell line (n = 18). Autonomic neurons derived from each FRDA iPSC line was first compared against those from the matched isogenic control iPSC line. A combined analysis was then performed across all three isogenic pairs, with individual of origin included as a covariate to account for inter-individual variability, allowing the effect of frataxin deficiency to be assessed independently of individual background (Online Resource 1). Individual of origin was the dominant source of transcriptional variation. Partitioning the variance of all 17,452 expressed genes attributed 42.1% to individual of origin, against 12.7% to disease (Fig. 4a). After removal of the individual of origin effect, the first principal component cleanly separated the FRDA samples from the isogenic control samples (Fig. 4b). The number of differentially expressed genes differed between iPSC lines (FA1, 28; FA2, 117; FA3, 6,597; FDR < 0.05).

**Fig. 4.**
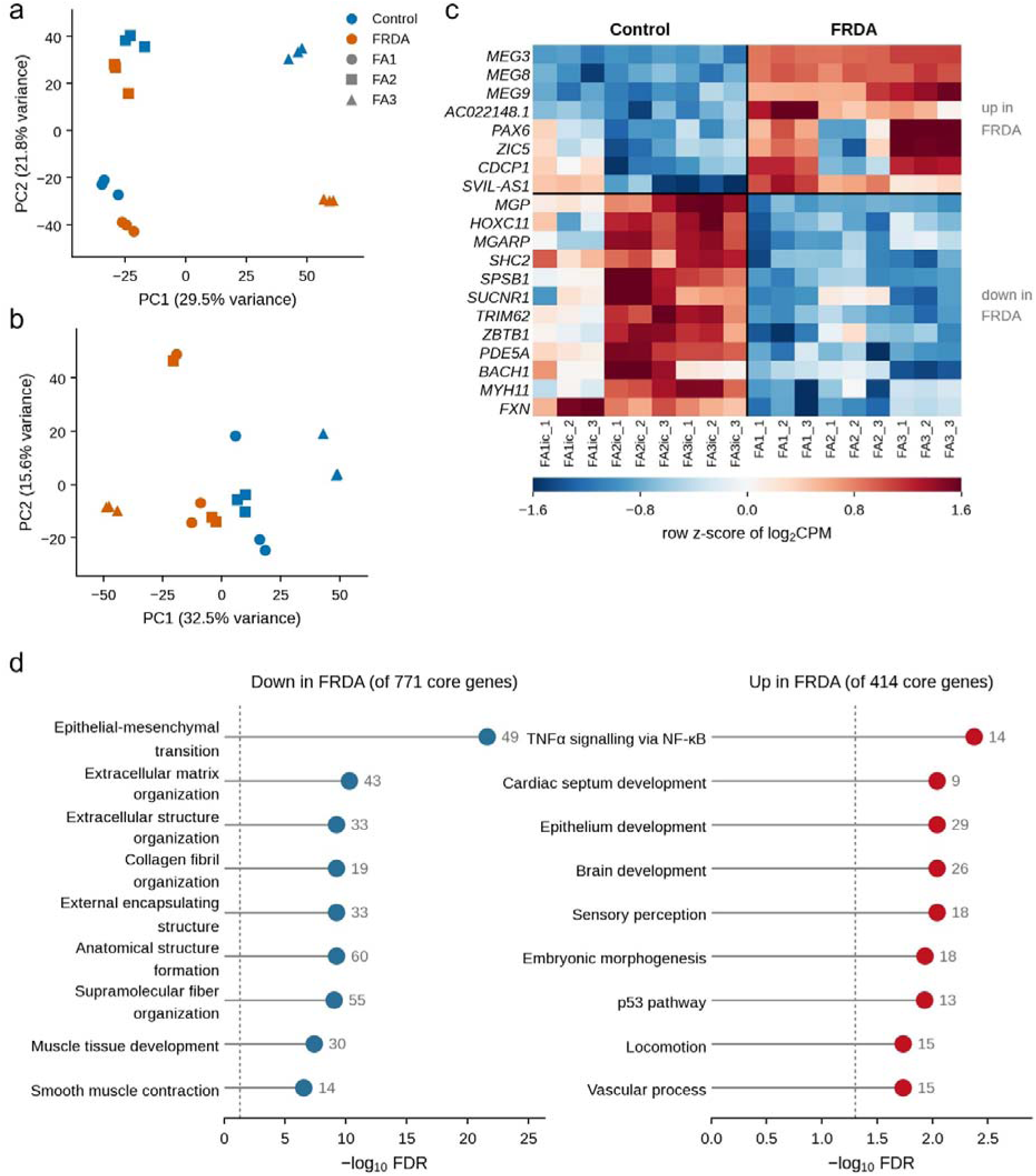
Transcriptomic analysis of autonomic neurons derived from Friedreich ataxia and isogenic control iPSCs. **a** Principal component analysis of the 2,000 most variable genes across all 18 samples; colour denotes disease status and symbol denotes patient line. **b** Principal component analysis after controlling for the patient-of-origin. **c** Heatmap of the 20 most significant conserved genes (12 down and 8 up in FRDA, ranked by FDR), shown as row *z*-scores of log_2_CPM. **d** Biological processes overrepresented among the 771 down-regulated and 414 up-regulated genes, tested against all 17,071 expressed genes (Fisher exact test, Benjamini–Hochberg FDR < 0.05); the nine most significant processes are shown for each direction and the number of genes in each process is given. n = 3 independent differentiations per line throughout

### Extracellular matrix programs are suppressed in FRDA autonomic neurons

A conserved core of 1,185 genes (414 up, 771 down in FRDA) was defined as genes differentially expressed overall, changed in the same direction in all three patient lines, and with a nominal *P* < 0.05 in at least two of the three lines (Fig. 4c). Overrepresentation analysis of this conserved core showed that down-regulated genes were strongly enriched for extracellular matrix, led by epithelial-mesenchymal transition (49 genes, FDR=2.7×10^-22^; e.g. *MGP*, *FN1*, *FSTL1*), extracellular matrix organisation (43 genes, FDR=5.2×10^-11^; e.g. *BGN*, *LTBP1*, *LAMB1*), and collagen fibril organisation (19 genes, FDR=6.1×10^-10^; e.g. *COL3A1*, *COL4A1*, *COL4A2*, *COL5A1*, *LOXL1*) (Fig. 4d). Beyond the terms shown in Fig. 4d, neuronal structural programs were also significantly suppressed, including neuron projection development (39 genes, FDR = 3.6 × 10^-3^; e.g. *MGARP*, *EFNA1*, *DMD*, *NREP*) and its positive regulation (16 genes, FDR = 2.0 × 10^-3^; e.g. *CUX2*, *VLDLR*, *DISC1*) (Online Resource 1). Up-regulated genes showed weaker enrichment overall, including stress response and developmental programmes, led by TNFα signalling via NF-κB (14 genes, FDR=4.2×10^-3^; e.g. *JUN*, *DUSP5*, *ID2*, *PDE4B*) and the p53 pathway (13 genes, FDR=1.2×10^-2^; e.g. *STEAP3*, *S100A4*, *IRAK1*) (Fig. 4d). All 449 significantly enriched terms and their constituent genes are given in Online Resource 1. Notably, the downregulation of extracellular matrix programs was a more dominant transcriptional feature of these autonomic neurons than the accompanying stress response.

### Top differentially expressed genes in FRDA autonomic neurons

The top 25 differentially expressed genes in FRDA iPSC derived autonomic neurons from three individuals with FRDA, ranked by adjusted p value, are listed in Table 1. Positive log fold change values indicate upregulation in FRDA iPSC derived autonomic neurons relative to their corrected isogenic controls. *MEG3* was the most significant differentially expressed gene overall (adjusted p = 1.43 × 10^-16^) and showed the largest magnitude change of any gene in the dataset, with a log fold change of 10.14, corresponding to marked upregulation in FRDA autonomic neurons. The imprinted locus transcripts *MEG8* and *MEG9* ranked second and third respectively. Downregulated genes among the top 25 included the matrix and adhesion associated genes *MGP*, *MGARP*. *FXN* was also among the top 25 downregulated genes (log_2_fold change = −1.04), confirming the expected loss of frataxin expression resulting from the GAA repeat expansion (Table 1).

**Table 1.** Top 25 differentially expressed genes in FRDA-iPSC-derived autonomic neurons from three individuals with FRDA, ranked by false discover rate (FDR). Positive log fold change values indicate upregulation in FRDA-iPSC-derived autonomic neurons relative to their isogenic controls

| Genes | FDR | Log<sub>2</sub> (fold-change) |
| --- | --- | --- |
| *MEG3* | 1.43e-16 | 10.14 |
| *MEG8* | 8.33e-11 | 4.23 |
| *MEG9* | 5.19e-09 | 3.86 |
| *AC022148.1* | 8.25e-06 | 2.87 |
| *HOXC11* | 8.54e-06 | -3.1 |
| *MGARP* | 8.54e-06 | -2.68 |
| *MGP* | 8.54e-06 | -4.47 |
| *SHC2* | 2.21e-05 | -0.94 |
| *SPSB1* | 2.56e-05 | -1.16 |
| *SUCNR1* | 3.31e-05 | -2.12 |
| *PAX6* | 6.32e-05 | 3.74 |
| *ZBTB1* | 7.61e-05 | -0.36 |
| *TRIM62* | 7.61e-05 | -0.97 |
| *BACH1* | 0.00011 | -0.81 |
| *PDE5A* | 0.00011 | -1.22 |
| *PPM1L* | 0.000124 | -1.01 |
| *ZIC5* | 0.000124 | 3.58 |
| *DEPP1* | 0.000124 | -1.22 |
| *MYH11* | 0.000124 | -2.38 |
| *CDCP1* | 0.000124 | 2.27 |
| *SVIL-AS1* | 0.000124 | 2.74 |
| *KCNA1* | 0.00016 | 3.46 |
| *MIR100HG* | 0.000167 | -0.57 |
| *FOXC1* | 0.000167 | -1.72 |
| *FXN* | 0.000167 | -1.04 |

### Maternally expressed transcripts at the imprinted 14q32 locus are de-repressed in FRDA autonomic neurons

The most strongly dysregulated transcript was the imprinted long non-coding RNA *MEG3*, elevated in all FRDA samples compared to the isogenic control samples (log_2_FC +10.14, FDR = 1.43 × 10^-16^; Fig. 5a). The neighbouring maternally expressed non-coding RNAs *MEG8* (log_2_FC +4.23) and *MEG9* (log_2_FC +3.86) were similarly elevated, whereas the paternally expressed protein-coding genes of the same *DLK1*-*DIO3* domain showed no consistent change (*DLK1*, FDR = 0.087; *DIO3*, FDR = 0.67); *RTL1* did not pass the expression filter and could not be assessed (Fig. 5a). This pattern is consistent with de-repression of the maternal allele, rather than activation of the whole 14q32 locus (Fig. 5b), and was reproduced by RT-qPCR in the same lines, with all three maternally expressed transcripts elevated and neither paternally expressed gene consistently changed (Fig. 5c).

**Fig. 5.**
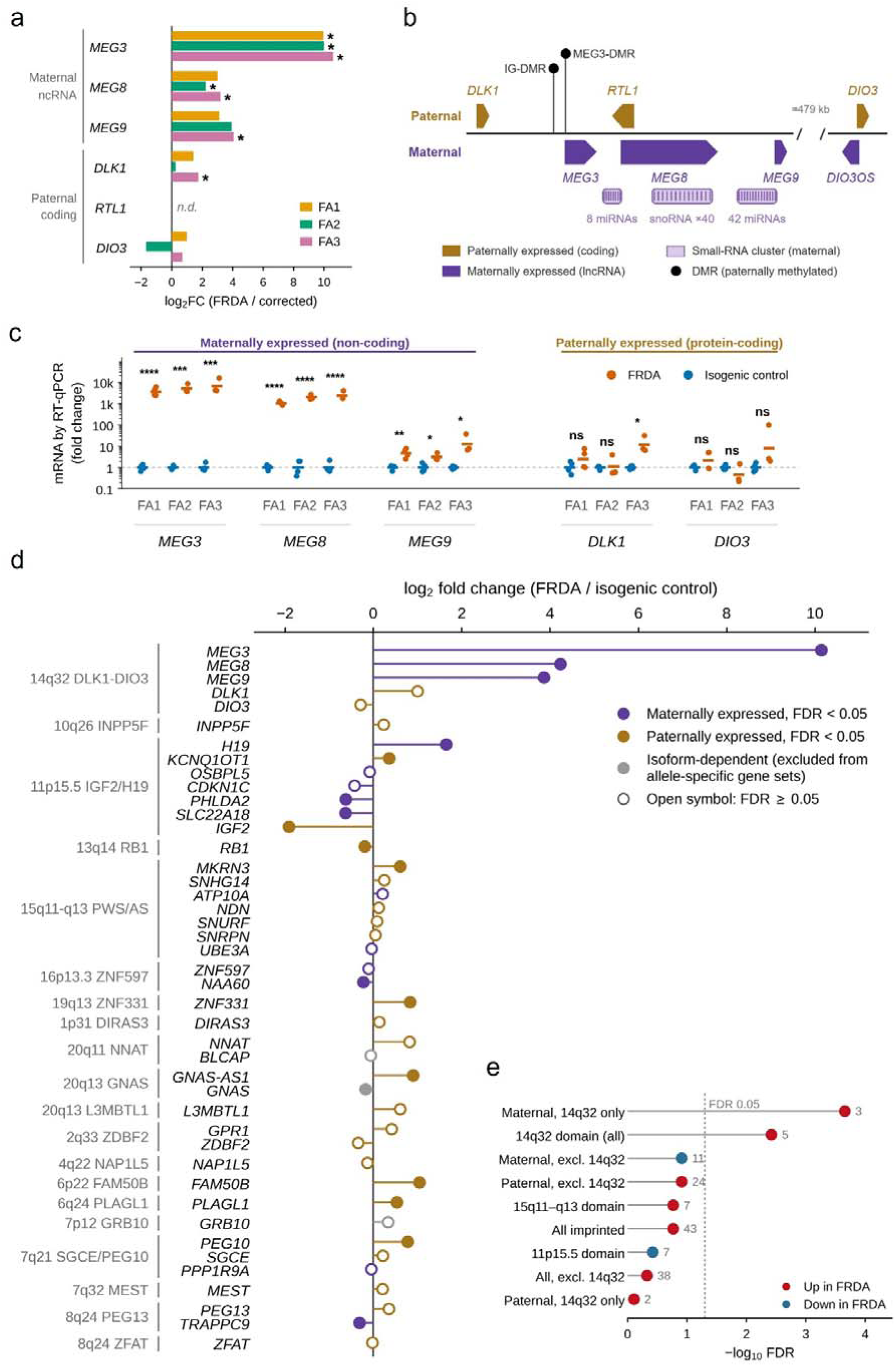
De-repression of maternally expressed transcripts at the 14q32 DLK1–DIO3 locus. **a** Per-line log_2_(fold change) for all 6 genes of the imprinted *DLK1–DIO3* domain at 14q32, grouped by parental allele of expression; *RTL1* was not detected above the expression filter (n.d.) and asterisks indicate FDR < 0.05 within that line. **b** Schematic of the 14q32 DLK1–DIO3 imprinted locus (chromosome 14, GRCh38). Gene positions are to scale; gene bodies are not. **c** RT-qPCR validation of the five quantifiable locus transcripts *MEG3*, *MEG8*, *MEG9*, *DLK1* and *DIO3*, normalised to *GAPDH* and expressed per patient relative to the matched isogenic corrected control (2^^−ΔΔCt^, log scale). *RTL1* was not quantified, due to overlapping sense and antisense transcripts. Points are independent samples; bars are geometric means. n = 3 independent differentiations. *p<0.05, **p < 0.01, ***p<0.001, ****p<0.0001 by unpaired Student’s t-test. **d** Log_2_ fold change (FRDA versus isogenic control) for 43 imprinted genes measurable in this dataset, spanning 20 imprinted domains, grouped by domain and coloured by the normally expressed parental allele. Filled symbols = FDR < 0.05; open symbols = FDR ≥ 0.05. Three genes that Geneimprint classifies as isoform-dependent (*GNAS*, *BLCAP*, *GRB10*) are shown in grey; these were tested individually but excluded from the allele-specific gene sets in **e**. Parental allele assignments are from the Geneimprint human catalogue. **e** Competitive gene-set tests (camera, limma) for nine imprinted gene sets: allele-specific sets inside and outside 14q32, whole-imprintome sets with and without 14q32, and the three domains with at least five measurable genes. Points show −log_10_ FDR after Benjamini–Hochberg correction across the nine sets tested; the number of genes in each set is given beside each point, and colour indicates the direction of change. Dashed line is FDR = 0.05. n = 3-4 independent differentiations per iPSC line

To determine whether this reflected a broader loss of imprinting, 43 imprinted genes across 20 imprinted domains were examined (Fig. 5d). Coordinated, direction-consistent dysregulation of a whole domain was restricted to 14q32 (FDR = 3.8 × 10^-3^) and was driven specifically by maternally expressed transcripts (FDR = 2.2 × 10^-4^). The paternally expressed genes of the 14q32 domain showed no coordinate change (FDR = 0.78) (Fig. 5e). Fifteen individual imprinted genes outside 14q32 were differentially expressed (including *H19*, *IGF2*, *PEG10* and *PLAGL1*) (Fig. 5d). However, neither of the other two domains with enough measurable genes to test (11p15.5 IGF2/H19 and 15q11–q13 PWS/AS) showed coordinated change (FDR > 0.05), and maternally expressed genes outside 14q32 showed a non-significant trend in the negative direction (FDR = 0.12) (Fig. 5e).

## Discussion

Autonomic dysfunction is a recognised but poorly understood feature of FRDA, and until now has lacked a human cellular model. By generating autonomic neurons from iPSCs of individuals with FRDA and their CRISPR/Cas9-corrected isogenic controls, this study provides the first controlled system to examine the cellular and molecular consequences of frataxin deficiency in this neuronal population. Frataxin-deficient autonomic neurons were hyperexcitable, with increased cell death, elevated mitochondrial reactive oxygen species, and mitochondrial fragmentation. Transcriptomic profiling revealed a convergent signature of extracellular matrix loss, alongside unexpected de-repression of the maternally imprinted genes in the 14q32 *DLK1-DIO3* locus, with the long noncoding gene *MEG3* the most significantly dysregulated gene across the dataset.

Clinical studies of autonomic function in FRDA have reached conflicting conclusions. Individuals with FRDA report substantial autonomic symptoms, with 80% reporting lower urinary tract symptoms, 64% bowel symptoms and 83% sexual symptoms [6, 25], and urinary dysfunction documented in 42.8% of 650 patients in the European registry [51]. Yet formal autonomic testing repeatedly fails to demonstrate autonomic failure. Valsalva responses, baroreflex sensitivity and sudomotor function are largely preserved, and the one consistent abnormality across studies is an elevated heart rate at rest and during orthostatic challenge [16, 18]. These observations do not point to denervation or the loss of autonomic neurons, but to altered autonomic output, consistent with the elevated plasma catecholamines reported in FRDA individuals [29, 39, 46]. In these individuals, such changes could equally reflect cardiac compensation, deconditioning, or reduced vagal tone [29], none of which can be distinguished clinically. Autonomic neurons in isolation carry none of these confounds, and their intrinsic excitability has not previously been measured in FRDA.

The present study shows that FRDA autonomic neurons fire more, at shorter interspike intervals, in longer and denser bursts, and did so in the absence of the influence of aberrant cardiac tissue or circulating factors. Hyperexcitability is therefore a cell-autonomous feature of FRDA autonomic neurons rather than secondary to cardiomyopathy. Burst frequency was unchanged while bursts became longer and denser, indicating altered intrinsic excitability rather than reorganisation of network activity. A similar neuronal phenotype has been described in iPSC-derived sympathetic neurons from individuals with long QT syndrome type 1, where functional hyperactivity accompanied sympathetically triggered arrhythmia [60], suggesting that sympathetic neuronal hyperexcitability may be a shared contributor to arrhythmic risk across distinct genetic cardiac diseases. Given that arrhythmia accounts for approximately one in eight cardiac related deaths in FRDA [38, 59], and since sympathetic drive is both pro-arrhythmic and pro-hypertrophic, these findings raise the possibility that autonomic nervous system dysfunction is a distinct and independent contributor to the pathogenesis of FRDA cardiomyopathy, upstream or parallel to the dysfunction seen in the better characterised cardiomyocytes.

The cellular and mitochondrial phenotype observed here in autonomic neurons mirrors observations made in other FRDA cell types and model systems. Increased cell death is well established in β-cells and neurons [15], dorsal root ganglion neurons [2, 56], and in FRDA iPSC-derived cardiomyocytes [30]. Elevated mitochondrial reactive oxygen species have been reported in mouse cerebellar granule neurons [1], primary rat ventricular myocytes [44], and FRDA iPSC-derived cardiomyocytes [4, 30]. A shift towards mitochondrial fragmentation has been described in frataxin-deficient yeast [31], fibroblasts from individuals with FRDA [22], and FRDA iPSC-derived cardiomyocytes [30]. Loss of mitochondrial membrane potential has been documented in FRDA mouse cerebellar granule neurons [1], dorsal root ganglion sensory neurons [2] and FRDA iPSC-derived cardiomyocytes [4], suggesting that frataxin deficiency leaves autonomic neurons metabolically vulnerable more broadly. High-frequency firing is metabolically expensive, so neurons firing more while carrying reduced energy reserves, common in FRDA, would be under chronic energetic strain, which may underlie the cell death observed. On that basis, reducing sympathetic firing could be useful as a clinical prophylaxis even without arrhythmia.

The phenotypic similarities noted above raises a broader discussion point. The same core phenotype recurs in frataxin-deficient yeast, primary fibroblasts and lymphoblasts from individuals with FRDA rodent cardiomyocytes and sensory neurons, mouse cerebellar neurons, iPSC-derived neurons and cardiomyocytes, and now human iPSC-derived autonomic neurons. The pathway linking frataxin deficiency to mitochondrial dysfunction therefore appears largely cell type independent, with lineage possibly setting the threshold at which the phenotype manifests rather than the mechanism itself. If so, cell types not yet examined would be expected to show similar dysfunction, for example, the enteric and pelvic neurons that innervate the organs implicated in the urinary, gastrointestinal and sexual symptoms reported in individuals with FRDA [6, 25, 51]. These neuronal populations have not yet been examined at a cellular level.

Individual of origin accounted for 42.1% of transcriptional variance, against 12.7% for disease, and DEG counts ranged from 28 to 6,597 per line. Analysis therefore focused on the signature consistent across all three iPSC lines (1,185 genes in total; 414 up, 771 down), and on *MEG3*, the most significantly dysregulated gene across the dataset. The dominant feature of that signature was coordinated loss of extracellular matrix programs, the strongest enrichment by several orders of magnitude, spanning fibrillar and basement membrane collagens, fibronectin and fibrillins, together with the enzymes that post-translationally mature collagen. Enzymes such as the lysyl hydroxylases *PLOD1* and *PLOD2* and the prolyl 3-hydroxylases *P3H2* and *P3H4*, are Fe(II)- and 2-oxoglutarate-dependent dioxygenases, which may be of particular interest given the iron dysmetabolism in FRDA [21, 48]. Loss of matrix and cell–substrate adhesion programs matter most in neurons that depend on long-range projection and stable target innervation. Consistent with this, neuron projection development was itself down-regulated (39 genes, FDR = 3.6 × 10^-3^) (Online Resource 1), and post-mortem studies describe loss of nerves and ganglia within the FRDA cardiac conduction system [19].

Beyond the extracellular matrix signature, the most significantly dysregulated gene across the dataset was the long noncoding RNA MEG3, which lies within the imprinted 14q32 DLK1–DIO3 domain. MEG3 was elevated approximately 1,000-fold, accompanied by the neighbouring noncoding RNAs MEG8 and MEG9. The effect was restricted to these maternally expressed transcripts. The paternally expressed protein-coding genes DLK1 and DIO3 of the same domain were not significantly altered, indicating maternal-allele de-repression rather than activation of the locus as a whole. Across the imprinted genes examined, individual genes were differentially expressed in several domains, but coordinated change at the level of a domain or parental allele was significant only for 14q32. This indicates a targeted effect at a single imprinted domain rather than a generalised loss of imprinting, and this allele-restricted pattern has not previously been described in FRDA. Aberrant methylation at the DLK1–DIO3 domain is a recognised consequence of reprogramming and extended culture [43, 57], so a clonal origin is plausible. However, 14q32 de-repression is among the most consistent findings in FRDA iPSC-derived cardiomyocytes, where MEG3 silencing reduced cell death [30]. It also recurs in independently derived isogenic FRDA lines from two other laboratories, in peripheral sensory neurons [26] and in neurons from three further donor backgrounds [41]. MEG3, MEG8 and MEG9 all rank within the top 30 genes of a pan-FRDA transcriptomic meta-analysis [36]. Given that FRDA pathophysiology involves repressive heterochromatin, dysregulation of an imprinted locus raises the possibility of a shared epigenetic mechanism. Whether 14q32 de-repression is a cause or a consequence of frataxin deficiency remains undetermined.

Frataxin deficiency produces a cell-autonomous phenotype in human iPSC-derived autonomic neurons, manifesting as hyperexcitability, mitochondrial fragmentation, oxidative stress, and increased cell death. The hyperexcitability provides the first direct cellular evidence for the increased sympathetic tone suggested by clinical studies of autonomic dysfunction in FRDA, and identifies the autonomic nervous system as a possible contributor to arrhythmic risk independent of cardiomyocyte pathology. Transcriptomic profiling identified co-ordinated de-repression of the maternally expressed transcripts of the imprinted 14q32 DLK1–DIO3 locus, led by *MEG3*, with the paternally expressed genes of the same locus unaffected and no comparable change at any other imprinted domain. Together, these findings establish iPSC-derived autonomic neurons as a useful model for a previously inaccessible aspect of FRDA, and position *MEG3* and the 14q32 locus as candidates for mechanistic and therapeutic investigation.

## Acknowledgments

The authors make no acknowledgments.

## Data Availability

All data supporting the findings of this study are available within the paper and its Supplementary Information. The Gene Expression Omnibus accession number for the RNA-seq dataset reported in this paper is GSE349562.

## Author contributions

J.G.L. and S.Y.L. conceived the project. J.G.L., M.D., A.P., L.C., M.D., M.N., and S.Y.L. were responsible for funding acquisition and experimental design. J.G.L., H.Z., L.L., S.W., J.C., and S.Y.L. performed experiments and data curation. J.G.L., H.Z., S.W., J.M.P., J.C., and S.Y.L. analyzed data. J.G.L., H.Z., A.P., M.D., L.C., M.D., D.M., M.N., and S.Y.L. interpreted results. J.G.L., H.Z., J.M.P., and S.Y.L. prepared figures. J.G.L. and S.Y.L. drafted manuscript. All authors participated in manuscript revision and approved the final version of the manuscript.

## Declarations

### Ethics approval and Consent to participate

The protocol for derivation of human primary fibroblasts was approved by the Institutional Review Board at the University of Alabama at Birmingham (IRB N131204003). All studies involving human material were performed in accordance with the ethical standards laid down in the 1964 Declaration of Helsinki and its later amendments. All participants gave written informed consent prior to inclusion in the study. No details that might disclose the identity of the participants are included.

### Consent to publish

Not applicable; no identifying information is included

## Online resources legends

**Online resource 1** Differential expression and enrichment analyses of autonomic neurons derived from three individuals with FRDA and their isogenic gene-corrected controls. Eight worksheets.

**OR1–OR3:** differential expression analysed separately for each patient line (OR1 = FA1, OR2 = FA2, OR3 = FA3), in each case comparing the FRDA line with its matched isogenic control; edgeR quasi-likelihood F tests within the ARMOR pipeline. All 17,452 genes retained by filterByExpr are listed with log□fold change, logCPM, F statistic, raw P and FDR; 28, 117 and 6,597 genes respectively reach FDR < 0.05.

**OR4:** differential expression for the combined analysis of all three patient lines, same pipeline.

**OR5:** overrepresentation analysis of the conserved core gene set against the Hallmark, Reactome and GO Biological Process collections. All 449 terms reaching FDR < 0.05 are listed (408 from down-regulated and 41 from up-regulated genes), each with its constituent genes.

**OR6:** genes reaching FDR < 0.05 in the analysis of all 18 samples with patient of origin included as a factor (2,876 genes).

**OR7:** the same analysis for all 17,452 genes tested, significant or not.

**OR8:** the conserved core of 1,185 genes, with per-line and combined statistics and the three selection criteria as columns.

